# Optimisation of Solid-Phase Antimicrobial Susceptibility Testing for Hydrophobic Antibiotics Using an Agarose-Based MIC Assay

**DOI:** 10.64898/2026.07.26.740829

**Authors:** Angela M. Kavanagh, Soumya Ramu, Gabrielle J. Lowe, Alison Hinton, Mark A. T. Blaskovich

## Abstract

The minimum inhibitory concentration (MIC) assay is the gold standard for evaluating antimicrobial activity^1^. However, conventional agar-based MIC methods often underestimate the potency of physicochemically complex compounds[1, 2]. Hydrophobic and adhesive molecules, such as lipoglycopeptide antibiotics, exhibit poor diffusion and non-specific binding to agar, leading to artificially elevated MIC values compared to broth-based methods[3]. This issue complicates accurate potency assessment and is particularly an issue when attempting resistance frequency (FOR) studies, which must be conducted on solid media.

Here, we developed a modified miniaturised agar MIC assay using 1% agarose, 0.002% Tween 80-supplemented tryptic soy broth (TSB), and a 24-well plate format[4]. These modifications improved compound dispersion, reduced matrix interactions, and lowered compound requirements. The optimised assay was validated with vancomycin, oritavancin, and dalbavancin against *Staphylococcus aureus* ATCC 43300 (MRSA) and *Streptococcus pneumoniae* ATCC 700677.

This efficient, cost-effective, high-throughput platform overcomes the limitations of traditional agar methods, enhancing reliability in evaluating challenging antimicrobials and supporting next-generation antibiotic development.

## Introduction

The minimum inhibitory concentration (MIC) assay remains the gold standard for evaluating the antimicrobial activity of compounds against bacterial isolates^1^. Traditionally performed using broth microdilution, this method provides a quantitative measure of growth inhibition in liquid culture and is widely regarded as the reference standard[5, 6]. Agar-based MIC assays serve as a valuable complementary approach, particularly for visualising inhibition zones and assessing antimicrobial effects under solid-phase conditions that more closely mimic certain environmental or clinical scenarios[5]. Furthermore, agar-based MIC methods are essential for frequency of resistance (FOR) assays[7], as they enable the quantification of resistant subpopulations capable of growing under defined antimicrobial selective pressures, providing important insights into the potential for resistance emergence[8].

However, conventional agar MIC methods can significantly underestimate the activity of certain compound classes, such as vancomycin derivatives. Highly hydrophobic or “sticky” molecules often exhibit strong surface interactions with the agar matrix, leading to poor diffusion, uneven distribution, and binding to agar components[4, 9]. These physicochemical limitations result in reduced apparent antimicrobial potency compared with results obtained in broth microdilution, even when compounds demonstrate clear and reproducible activity in liquid media[10]. Such discrepancies highlight the need for assay adaptations tailored to the specific properties of the test compounds.

In the present study, we encountered this challenge while evaluating a series of highly adhesive compounds that showed potent inhibition in broth microdilution but little to no activity under standard agar conditions. To overcome diffusion and surface-binding issues, we modified key parameters of the traditional agar MIC assay. These adaptations enabled a more reliable assessment of antimicrobial potential on solid media and underscore the importance of optimising assay design when working with physicochemically complex molecules.

## Results

In the comparative evaluations of broth microdilution and agar-based assays testing glycopeptide derivatives, vancomycin was the only compound that retained consistent antimicrobial activity across all tested conditions, exhibiting an MIC of 1 µg/mL. In contrast, the more lipophilic agents, such as oritavancin, dalbavancin, and our novel ‘vancapticin’ vancomycin derivative compounds[11], displayed evident medium-dependent differences. In both standard broth microdilution and the optimized miniaturized TSB–1% agarose–0.002% Tween 80 assay, these compounds demonstrated potent activity with MIC values ranging from 0.03 to 0.06 µg/mL. However, under conventional TSA conditions, oritavancin MICs increased to 4 µg/mL, representing an approximately 67-fold reduction in apparent potency. Dalbavancin showed a more modest yet still significant 8.3-fold reduction in activity on TSA, while the vancomycin derivatives exhibited a similar trend (Table 1). These observations are consistent with previous reports highlighting the challenges of assessing hydrophobic or amphiphilic molecules in standard agar matrices due to poor diffusion and non-specific binding[4, 10].

**Table 1.** MIC result Broth vs Agarose vs TSA.

| Compound<br>Name | *Staphylococcus aureus*<br>ATCC 43300 |  |  | *Streptococcus pneumoniae*<br>ATCC 700677 |  |  |
| --- | --- | --- | --- | --- | --- | --- |
|  | MHB Broth | Tryptic soy<br>agar (TSA) | Modified<br>agar: Tryptic<br>soy broth with<br>1% agarose +<br>0.002% Tween) | MHB Broth | Tryptic soy<br>agar (TSA) | Modified<br>agar: Tryptic<br>soy broth with<br>1% agarose +<br>0.002% Tween) |
| Vancomycin | 1/0.5 | 2 | 1 | 1 | 2 | 2 |
| Oritavancin | 0.06/0.003 | 1/4 | 0.03/0.015 | 0.06 | 4 | 0.06 |
| Dalbavancin | 0.006/≤0.003 | 0.5/0.25 | 0.03/0.015 | 0.03/0.015 | 0.5/0.25 | 0.06 |
| MCC_00080 | 0.06/0.03 | 0.25 | 0.06 | 0.125 | 0.5 | 0.125/0.06 |
| MCC_05145 | 0.006/≤0.003 | >1 | 0.06 | 0.06/0.03 | >1 | 0.06 |

These initial findings validated the utility of the modified agar assay and enabled the implementation of frequency-of-resistance (FOR) assays under optimised conditions. The assays produced highly reproducible results, with stable titers of 18-hour stationary-phase cultures reliably determined (Table 2). This allowed accurate quantification of spontaneous resistance frequencies for *Staphylococcus aureus* ATCC 43300 (MRSA) and *Streptococcus pneumoniae* ATCC 700677 at concentrations up to 16× MIC (Table 3). Such low resistance frequencies are consistent with the multi-target mechanisms of action reported for lipoglycopeptides.

**Table 2.** Titer of 18-hour stationary phase culture.

| *Staphylococcus aureus* ATCC 43300 |  |  |  |  |  |
| --- | --- | --- | --- | --- | --- |
| Dilution<br>Tube | CFU/plate |  | Average<br>CFU/plate | Dilution<br>factor | Average<br>CFU/mL |
| 10<sup>-6</sup> | 354 | 349 | 351.5 | 1.00x10<sup>6</sup> | 7.03x10<sup>9</sup> |
| 10<sup>-7</sup> | 418 | 302 | 360 | 1.00x10<sup>7</sup> | 7.20x10<sup>10</sup> |
| 10<sup>-8</sup> | 175 | 152 | 163.5 | 1.00x10<sup>8</sup> | 3.27x10<sup>11</sup> |
|  |  |  |  |  | 1.35x10<sup>11</sup> |
| *Streptococcus pneumoniae* ATCC 700677 |  |  |  |  |  |
| Dilution<br>Tube | CFU/plate | Average<br>CFU/plate | Dilution<br>factor | Average<br>CFU/mL | Dilution<br>Tube |
| 10<sup>-6</sup> | 414 | 474 | 444 | 1.00x10<sup>6</sup> | 8.88x10<sup>9</sup> |
| 10<sup>-7</sup> | 58 | 79 | 68.5 | 1.00x10<sup>7</sup> | 1.37x10<sup>10</sup> |
| 10<sup>-8</sup> | 6 | 5 | 5.5 | 1.00x10<sup>8</sup> | 1.10x10<sup>10</sup> |
|  |  |  |  |  | 1.12x10<sup>10</sup> |

**Table 3.**
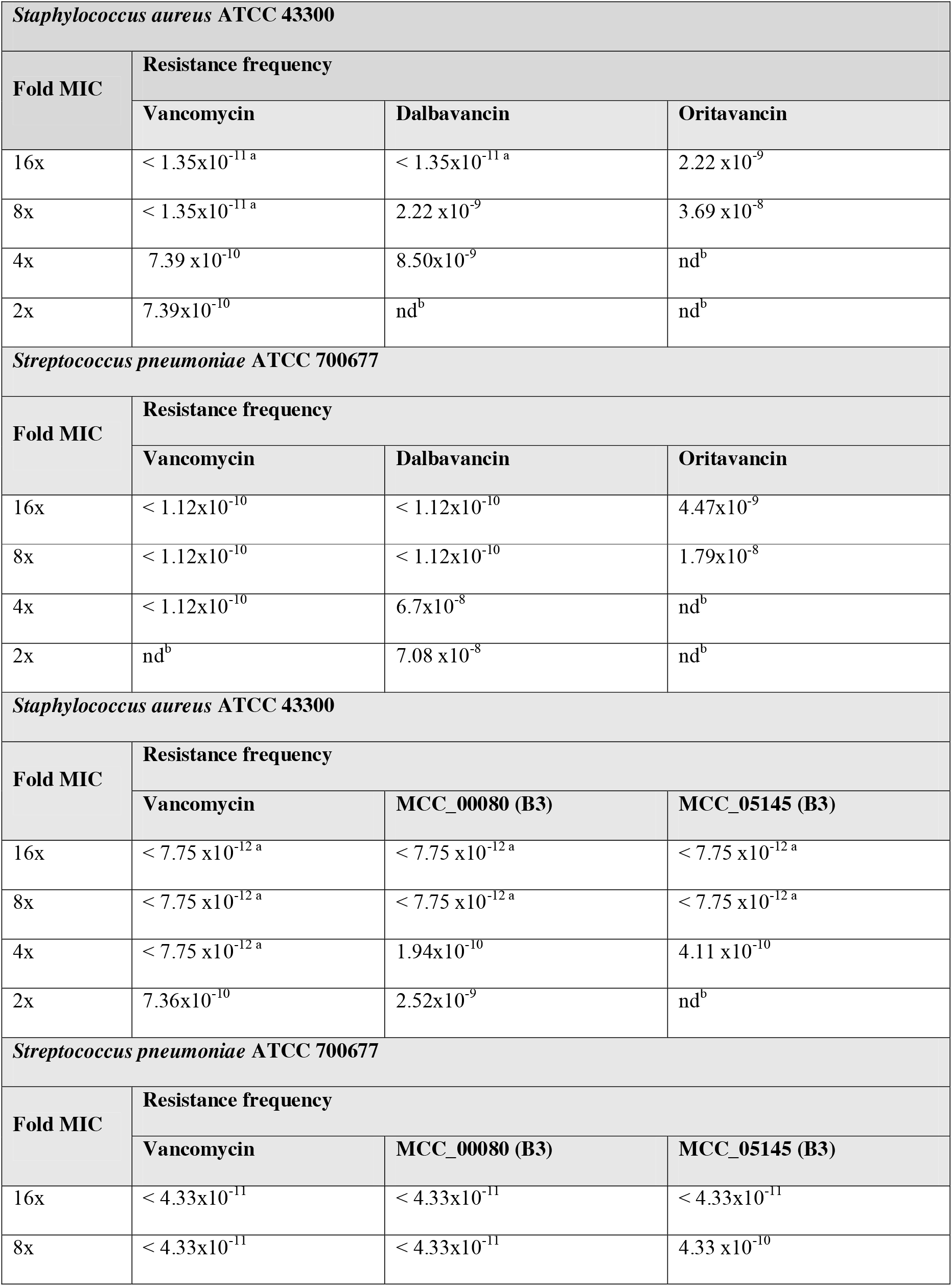
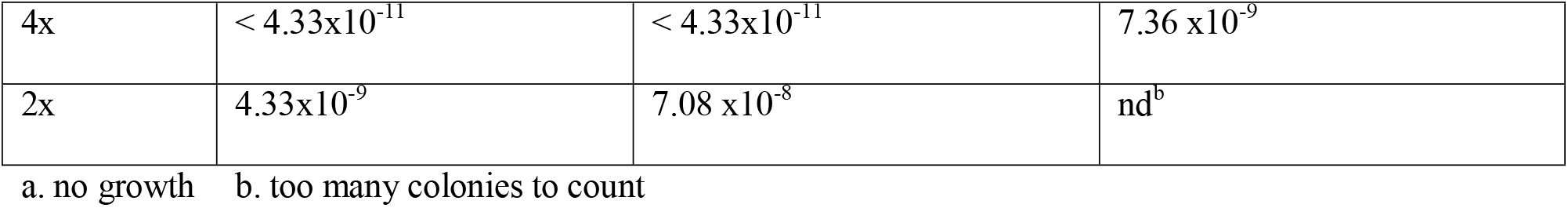
Resistance Frequency Test Results using modified conditions.

| *Staphylococcus aureus* ATCC 43300 |  |  |  |
| --- | --- | --- | --- |
| Fold MIC | Resistance frequency |  |  |
|  | Vancomycin | Dalbavancin | Oritavancin |
| 16x | $< 1.35 \times 10^{-11}$ a | $< 1.35 \times 10^{-11}$ a | $2.22 \times 10^{-9}$ |
| 8x | $< 1.35 \times 10^{-11}$ a | $2.22 \times 10^{-9}$ | $3.69 \times 10^{-8}$ |
| 4x | $7.39 \times 10^{-10}$ | $8.50 \times 10^{-9}$ | nd<sup>b</sup> |
| 2x | $7.39 \times 10^{-10}$ | nd<sup>b</sup> | nd<sup>b</sup> |
| *Streptococcus pneumoniae* ATCC 700677 |  |  |  |
| Fold MIC | Resistance frequency |  |  |
|  | Vancomycin | Dalbavancin | Oritavancin |
| 16x | $< 1.12 \times 10^{-10}$ | $< 1.12 \times 10^{-10}$ | $4.47 \times 10^{-9}$ |
| 8x | $< 1.12 \times 10^{-10}$ | $< 1.12 \times 10^{-10}$ | $1.79 \times 10^{-8}$ |
| 4x | $< 1.12 \times 10^{-10}$ | $6.7 \times 10^{-8}$ | nd<sup>b</sup> |
| 2x | nd<sup>b</sup> | $7.08 \times 10^{-8}$ | nd<sup>b</sup> |
| *Staphylococcus aureus* ATCC 43300 |  |  |  |
| Fold MIC | Resistance frequency |  |  |
|  | Vancomycin | MCC_00080 (B3) | MCC_05145 (B3) |
| 16x | $< 7.75 \times 10^{-12}$ a | $< 7.75 \times 10^{-12}$ a | $< 7.75 \times 10^{-12}$ a |
| 8x | $< 7.75 \times 10^{-12}$ a | $< 7.75 \times 10^{-12}$ a | $< 7.75 \times 10^{-12}$ a |
| 4x | $< 7.75 \times 10^{-12}$ a | $1.94 \times 10^{-10}$ | $4.11 \times 10^{-10}$ |
| 2x | $7.36 \times 10^{-10}$ | $2.52 \times 10^{-9}$ | nd<sup>b</sup> |
| *Streptococcus pneumoniae* ATCC 700677 |  |  |  |
| Fold MIC | Resistance frequency |  |  |
|  | Vancomycin | MCC_00080 (B3) | MCC_05145 (B3) |
| 16x | $< 4.33 \times 10^{-11}$ | $< 4.33 \times 10^{-11}$ | $< 4.33 \times 10^{-11}$ |
| 8x | $< 4.33 \times 10^{-11}$ | $< 4.33 \times 10^{-11}$ | $4.33 \times 10^{-10}$ |
| 4x | $< 4.33 \times 10^{-11}$ | $< 4.33 \times 10^{-11}$ | $7.36 \times 10^{-9}$ |
| 2x | $4.33 \times 10^{-9}$ | $7.08 \times 10^{-8}$ | nd<sup>b</sup> |

a. no growth b. too many colonies to count

## Discussion

The miniaturised agar MIC assay developed in this study, incorporating 1% agarose, 0.002% Tween 80 in TSB, and a 24-well plate format, successfully enabled the evaluation of hydrophobic lipoglycopeptide antibiotics (oritavancin and dalbavancin) that typically exhibit poor performance in conventional agar-based systems. All three compounds demonstrated potent activity, with vancomycin MICs of 1–2 µg/mL against both *S. aureus* ATCC 43300 (MRSA) and *S. pneumoniae* ATCC 700677, consistent with established susceptibility profiles. Notably, oritavancin and dalbavancin exhibited markedly lower MIC values (0.015– µg/mL), highlighting their superior potency against these Gram-positive pathogens under the optimised solid phase conditions. These results closely align with broth microdilution data, confirming that the modifications effectively mitigate the diffusion and binding limitations previously observed with standard agar media for these “sticky” molecules.

The inclusion of a low concentration of Tween 80 (0.002%) as a non-ionic surfactant, combined with agarose rather than bacteriological agar, likely improved compound solubility and diffusion while minimising non-specific binding to the matrix. This approach addresses a well-documented challenge in antimicrobial susceptibility testing of lipophilic agents, where traditional agar methods can lead to falsely elevated MICs[4].The miniaturised format also offers practical advantages, including reduced compound consumption, higher throughput, and easier visual readout, making it particularly suitable for early-stage screening of physiochemically challenging compounds[9].

Resistance frequency determinations further supported the therapeutic potential of these agents. Spontaneous resistance frequencies to vancomycin were extremely low or undetectable at 2–8× MIC, consistent with its established clinical profile. For the more potent lipoglycopeptides, resistance frequencies remained low overall (generally in the range of 10□□ to 10□□), even at concentrations up to 32× MIC. While a small number of colonies were observed for dalbavancin and oritavancin at 4–16× MIC, the calculated frequencies indicate a relatively high genetic barrier to resistance. These findings are encouraging, as low resistance emergence is a critical attribute for next generation antibiotics targeting multidrug-resistant Gram-positive bacteria[10, 12].

The low resistance frequencies observed for MCC_00080 and MCC_05145 are particularly significant given their enhanced potency. Lipoglycopeptides such as oritavancin and dalbavancin typically exhibit low spontaneous mutation rates, attributed to their multiple mechanisms of action (including membrane disruption in addition to cell wall inhibition). The current results align with this profile and suggest that the test compounds maintain this advantageous property. The modified agar assay was critical in enabling accurate assessment of these hydrophobic molecules, as standard agar methods often underestimate their true activity due to poor diffusion and matrix binding.

It is important to acknowledge certain limitations of the current study. The resistance frequency assays were performed using a single strain of each species, and results may vary with more diverse clinical isolates. Additionally, while the miniaturised agar assay performed well for these lipoglycopeptides, further validation with a broader panel of hydrophobic compounds and additional bacterial species is warranted. Colony counting at higher compound concentrations was occasionally complicated by growth patterns (e.g., streaking), which may introduce minor variability in frequency calculations.

In conclusion, the modified miniaturised agar MIC assay provides a robust, reliable and resource-efficient platform for assessing the antimicrobial activity of hydrophobic compounds on solid media. By overcoming the diffusion and binding issues inherent to conventional agar methods, this approach bridges the gap between broth and agar susceptibility testing. The potent MIC values and low resistance frequencies observed for oritavancin and dalbavancin reinforce their value as important therapeutic options and demonstrate the utility of tailored assay design in antimicrobial drug development.

## Materials and Methods

### Miniaturised agar MIC Assay

Tested compounds were added to Tryptic Soy Broth (BD, Cat. No, 211825), 1% Agarose, 0.002% Tween-80 (TSB-AT) in 15 mL Falcon tubes to achieve a final two-fold dilution range from 4 – 0.008 µg/mL. Compound-containing agar was aliquoted in duplicate, 1 mL per well, into 24- non-TC plates (PS; Corning, Cat. No. 3738).

Bacteria were cultured in Mueller-Hinton broth (MHB; BD, Cat. No. 211443) at 37 ºC overnight, then diluted 40-fold and incubated at 37 ºC for a further 2-3 hours. The resultant mid-log phase cultures were serially diluted in MHB from 10^9^ CFU/mL to give a final cell density of 10^6^ CFU/mL^1^. 10 µL of inoculated media was dispensed onto the surface of the agar in each well, except for the negative growth control wells, to which 10 µL of uninoculated media was added. The plates were covered and incubated at 37 ºC for 20 hours.

Inhibition of bacterial growth was determined visually, where the MIC was recorded as the lowest compound concentration with no visible growth.

### Resistance frequency assay

To determine the titer of the bacteria tested in an 18 h stationary-phase culture, each strain was cultured in MHB at 37 ºC overnight. Then, serially diluted to a 10^-8^ dilution. The dilutions 10^−6^, 10^−7^, and 10^−8^ were used[7]. 50 μL of these dilutions were added to compound-free petri plates containing Tryptic Soy Broth (TSB; BD, Cat No. 211825) with 1% agarose and 0.002% Tween80, plated in duplicate[3]. Plates were incubated overnight at 37°C.

Each tested compound was added to Tryptic Soy Broth with 1% agarose, 0.002% Tween80 in a glass bottle maintained at 45-50 °C in a water bath, giving a final concentration 4X (vancomycin) / 32X (oritavancin and dalbavancin) above MIC. The compound-containing mixture was added to petri dishes, giving a final volume of 20 mL per plate. The remaining agarose-containing compound mixture was then serially diluted two-fold, and the process was repeated until 1X / 4X / 8X MIC values were reached. For a neat culture, 10 µL was plated, for a 10-1 culture, 200 µL was plated onto the compound containing agarose and spread out evenly across the plate.

Plates were incubated overnight at 37°C for 20 hours.

### Detection and analysis

Colony counts were completed after 24 hours of incubation. The well-separated visible colonies were counted; any colonies that grew together were not included in the count.

#### Resistance Frequency Determination

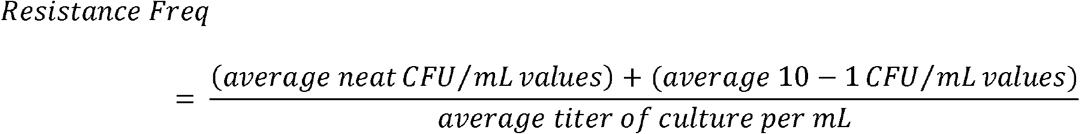

## Acknowledgement

We thank National Health and Medical Research Council (NHMRC) Principal Research Fellow (APP1059354) and former Australia Fellow (AF511105). J.A.R. is a NHMRC Career Development Fellow (APP1048652). This work was supported by a Wellcome Trust Seeding Drug Discovery Award 094977/Z/10/Z, and NHMRC Project Grants APP631632 and APP1026922. R.A.G.S acknowledges the support of the UK Medical Research Council and the UK NIHR Biomedical Research Council at Guy’s Hospital, London. The following strains were provided by the Network on Antimicrobial Resistance. Strains S. aureus ATCC 4300 and S. pneumoniae ATCC 700677 were acquired from American Type Culture Collection (ATCC).

## Notes

### Competing Interest Statement

The authors have declared no competing interest.

